# Using dual-calibrated functional MRI to map brain oxygen supply and consumption in multiple sclerosis

**DOI:** 10.1101/2021.01.07.425819

**Authors:** Hannah L Chandler, Rachael C Stickland, Michael Germuska, Eleonora Patitucci, Catherine Foster, Shona Bhome-Dhaliwal, Thomas M Lancaster, Neeraj Saxena, Sharmila Khot, Valentina Tomassini, Richard G Wise

**Affiliations:** CUBRIC, School of Psychology, Cardiff University, Cardiff, United Kingdom; Department of Physical Therapy and Human Movement Sciences, Northwestern University, Chicago, IL, USA; Wales Institute of Social and Economic Research and Data, Cardiff University, Cardiff; Cardiff University School of Medicine, Cardiff; Department of Psychology, University of Bath, Bath, UK; Department of Anaesthetics, Intensive Care and Pain Medicine, Cwm Taf Morgannwg, University Health Board, Abercynon, UK; Institute for Advanced Biomedical Technologies, Department of Neuroscience, Imaging and Clinical Sciences, University G. d’Annunzio of Chieti-Pescara, Chieti, Italy; MS Centre, Neurology Unit, “SS. Annunziata” University Hospital, Chieti, Italy; Division of Psychological Medicine and Clinical Neurosciences, School of Medicine, Cardiff University, Cardiff, UK; Helen Durham Centre for Neuroinflammation, University Hospital of Wales, Cardiff, UK

**Author notes:** Equal contribution (first author) (Hannah L Chandler, Rachael C Stickland). Equal contribution (senior author) (Valentina Tomassini, Richard G Wise). **Corresponding author** Professor Richard Wise, Institute for Advanced Biomedical Technologies (ITAB), Department of Neuroscience, Imaging and Clinical Sciences, University G. D’Annunzio of Chieti-Pescara, Chieti, Italy.

**Keywords:** MS, damage, disability, dual-calibrated functional MRI, oxygen, perfusion

## Abstract

Evidence suggests that cerebrovascular function and oxygen consumption are altered in multiple sclerosis (MS). Here, we quantified the vascular and oxygen metabolic MRI burden in patients with MS (PwMS) and assessed the relationship between these MRI measures of and metrics of damage and disability. In PwMS and in matched healthy volunteers, we applied a newly developed dual-calibrated fMRI method of acquisition and analysis to map grey matter (GM) cerebral blood flow (CBF), oxygen extraction fraction (OEF), cerebral metabolic rate of oxygen consumption (CMRO_2_) and effective oxygen diffusivity of the capillary network (D_C_). We also quantified physical and cognitive function in PwMS and controls. There was no significant difference in GM volume between 22 PwMS and 20 healthy controls (*p*=0.302). Significant differences in CBF (PwMS vs. controls: 44.91 ± 6.10 vs. 48.90 ± 5.87 ml/100g/min, *p*=0.010), CMRO_2_ (117.69 ± 17.31 vs. 136.49 ± 14.48 μmol/100g/min *p*<0.001) and D_C_ (2.70 ± 0.51 vs. 3.18 ± 0.41 μmol/100g/mmHg/min, *p*=0.002) were observed in the PwMS. No significant between-group differences were observed for OEF (PwMS vs. controls: 0.38 ± 0.09 vs. 0.39 ± 0.02, *p*=0.358). Regional analysis showed widespread reductions in CMRO_2_ and D_C_ for PwMS compared to healthy volunteers. There was a significant correlation between physiological measures and T2 lesion volume, but no association with current clinical disability. Our findings demonstrate concurrent reductions in oxygen supply and consumption in the absence of an alteration in oxygen extraction that may be indicative of a reduced demand for oxygen (O_2_), an impaired transfer of O_2_ from capillaries to mitochondria, and/or a reduced ability to utilise O_2_ that is available at the mitochondria. With no between-group differences in GM volume, our results suggest that changes in brain physiology may precede MRI-detectable GM loss and thus may be one of the pathological drivers of neurodegeneration and disease progression.

## Introduction

Evidence suggests that brain physiology is altered in multiple sclerosis (MS) (Paling, Golay, Wheeler-Kingshott, Kapoor, & Miller, 2011), where reductions in cerebral perfusion precede lesion formation (Wuerfel et al., 2004) and can occur in the absence of significant tissue volume loss (Debernard et al., 2014). Consistently, changes in cerebrovascular reactivity (CVR) occur in MS, correlating with the degree of damage (Marshall, Chawla, Lu, Pape, & Ge, 2016; Marshall et al., 2014). In parallel with changes in brain energy supply, reductions in brain energy consumption have been detected using [^18^F] FDG and [^15^O] PET (Paulesu et al., 1996), (Brooks et al., 1984; Sun, Tanaka, Kondo, Hirai, & Ishihara, 1994; Sun, Tanaka, Kondo, Okamoto, & Hirai, 1998). These changes overall suggest a neurovascular unit malfunction that accompanies an altered capacity for energy utilisation in the MS brain. They also suggest that patients with MS may experience hypoxic-like tissue states that are linked to inflammatory processes and involve a cascade of events that may chronically lead to further tissue damage (Davies et al., 2013; Haider et al., 2016; Law et al., 2004; Trapp & Stys, 2009; Yang & Dunn, 2015) and disability (Ge et al., 2012) (West et al., 2020). The relationship between altered brain oxygen supply and consumption in MS, however, remains to be elucidated. Additional information is required to identify whether reduced vascular function may be limiting energy consumption or whether reduced vascular performance is secondary to reduced energy demand or a reduced ability of the tissue to access and use metabolic substrates.

The development of novel multi-parametric MRI methods that map relevant physiological parameters regionally across the cortex may provide insight into the disease processes that contribute to tissue damage and disability. Studies mapping the effects of MS on brain energy consumption in humans have so far been limited to PET. Recently, non-invasive MRI-based measurements of venous oxygenation have become available that offer a global or regional estimate of OEF, depending on the chosen vein (Fan et al., 2015b; Ge et al., 2012). However, the territory drained by the sampled vein(s) remains uncertain and so localisation of OEF and, therefore, CMRO_2_, once CBF estimates are combined with OEF, are not feasible. Dual-calibrated fMRI (dc-fMRI) methods (Bulte et al., 2012; Gauthier & Hoge, 2013; Germuska & Wise, 2019; Wise, Harris, Stone, & Murphy, 2013) have demonstrated their ability to map CBF, OEF and CMRO_2_ in GM (Merola, Germuska, Murphy, & Wise, 2018) in clinical cohorts (Hoge, 2012) and with pharmacological intervention (Merola et al., 2017). Arterial spin labelling (ASL), combined with a biophysical model of the MR signals, permits a regional mapping of CBF, OEF and absolute CMRO_2_ in GM (Germuska et al., 2019; Germuska et al., 2016), as well as the estimation of effective oxygen diffusivity of the capillary network (D_C_) (Germuska et al., 2019).

In this study we use dc-fMRI to quantify in CBF, OEF, CMRO_2_, and D_C_ across the GM of patients with relapsing-remitting MS (PwMS) and test for differences with age and sex matched healthy controls, under the hypothesis of a generalised vascular and metabolic dysregulation in the GM of the MS brain. With the addition of D_C_, we extended our investigation to include the function of the GM capillary network that offers the opportunity to improve our understanding of the pathophysiological mechanisms of tissue dysfunction and damage. We further investigated the relationship between measures of vascular and oxygen metabolic function and metrics of brain damage and disability. To reduce the effect of GM loss that may bias the cerebral physiological parameters, we performed analysis restricting our estimate of those physiological parameters to voxels with a high proportion of GM.

## Methods

### Participants

We recruited PwMS and age and sex matched healthy controls of mixed handedness. All patients were recruited at the University Hospital of Wales in Cardiff, UK. All PwMS had a diagnosis of MS (Thompson et al., 2018) with a relapsing-remitting course (Lublin et al., 2014). Eligibility criteria for PwMS included no change in medication and no relapse in the 3 months prior to study entry. Participants has no history (within the preceding 2 years) of, or were being treated for, any significant cardiac or respiratory, and were not regular smokers within the previous 6 months. Written consent was obtained according to the protocol approved by the NHS Research Ethics Committee, Wales, UK.

### Data collection

#### Clinical data and behavioural assessments

All participants completed a socio-demographic and lifestyle questionnaire. Tests from the MS Functional Composite (Cutter et al., 1999) were carried out on all participants: 9-Hole Peg Test (9-HPT) for arm/hand function, Timed 25-Foot Walk (T25-FW) for leg function/ambulation, the Paced Auditory Serial Addition Test (PASAT) 2 and 3 seconds as a measure of cognitive function. The Symbol Digit Modalities Test (SDMT) was also used to assess cognitive function (Benedict et al., 2017). Visual acuity was assessed, in each eye separately, with a SLOAN letter chart (*Precision Vision*) at 100% contrast and scored in two ways: (1) the smallest letter size (in M-units) where 3 out of 5 letters were correctly read, and (2) a cumulative score of total letters read, out of 60. Participants were tested with corrected vision. For patients only, disability and disease impact were assessed with the Multiple Sclerosis Impact Scale (MSIS-29) (Hobart, Lamping, Fitzpatrick, Riazi, & Thompson, 2001) and the Fatigue Scale for Motor and Cognitive Functions (FSMC) (Penner et al., 2009). Clinical records and a short interview on disease history and impact gave information on: disease onset, Expanded Disability Status Scale (EDSS) score (Kurtzke, 1983), relapse history, experience of pain on an average day (rating from 0 being no pain to 10 being worst pain imagined), and impact of MS on occupation.

### MRI acquisitions

All MRI data were acquired on Siemens Prisma 3T MRI scanner (*Siemens Healthineers, Erlangen, Germany*), using a 32-channel receive-only head coil.

#### Structural scans

A magnetization-prepared rapid acquisition with gradient echo (MPRAGE) T1-weighted scan was acquired for registration and brain segmentation purposes (matrix 165 × 203 × 197, 1mm^3^ resolution, TR/TE = 2100/3.24ms). A 3D T2-weighted Fluid Attenuated Inversion Recovery (FLAIR) image (1mm^3^ isotropic resolution, 256 slices, TR/TE = 5000/388ms) and T2/Proton Density dual-echo image (in-plane resolution 0.8 × 0.8 mm, 41 (3.9 mm thick) slices TR/TE1/TE2= 4050/11/90ms) were acquired for lesion identification.

#### Quantitative functional scans

We used a home-written dual-excitation (Schmithorst et al., 2014) PCASL-based dual-calibration method with background suppression and pre-saturation (Germuska & Wise, 2019; Okell, Chappell, Kelly, & Jezzard, 2013) that acquire CBF and BOLD signals together to map changes in oxygen consumption and cerebrovascular function across GM. Scan parameters were: TE1: 10ms, TE2: 30ms, TR: 4400ms, in-plane resolution 3.4 × 3.4 mm, 16 (7mm thick) slices with 20% slice gap and GRAPPA acceleration factor 3 (**Figure 1**). The acquisition time for the PCASL sequence was 18 minutes, during which interleaved periods of hypercapnic and hyperoxic gases were delivered (Germuska et al., 2016; Germuska & Wise, 2019). Hypercapnia blocks involved delivering 5% CO_2_, whereas for the hyperoxia blocks, 50% O_2_ was delivered with short periods of 100% O_2_ at the start and 10% O_2_ at the end delivered to accelerate the transitions to the hyperoxic state and back to baseline, respectively. Physiological monitoring was conducted throughout measurement of the heart rate (HR) and O_2_ (%) saturation. Partial pressure of end tidal O_2_ (P_ET_O_2_) and CO_2_ (P_ET_CO_2_) were collected from the participant’s facemask and recorded using gas analyzer software from AD Instruments (PowerLab^®^, ADInstruments, Sydney, Australia). A blood sample was drawn via a finger pick with a 1.8mm lancet, and analysed with the HemoCue^®^ Hb 301+ system in accordance with the manufacturer’s guidelines (*Hemo Ängelholm, Sweden*). This gave a reading of blood haemoglobin concentration ([Hb]), in grams per litre (g/L), for use in the calculation of OEF. This was performed within the hour before the start of the MRI scan.

**Figure 1.**
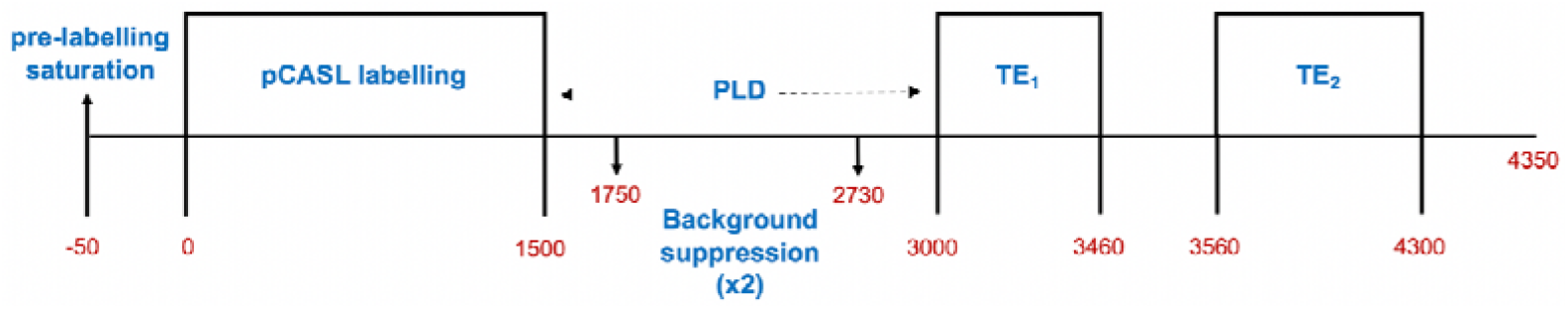
Graphical representation of the sequence parameters and timings (ms) for dc-fMRI dual-excitation sequence. TE_1_ represents the block of slices acquired for ASL weighted signal, whereas TE_2_ represents the block of slices acquired for BOLD weighted signal. *Abbreviations:* Blood oxygen level dependent (BOLD), arterial spin labelling (ASL), cerebral blood flow (CBF), oxygen extraction fraction (OEF), cerebral metabolic rate of oxygen consumption (CMRO_2_) and capillary oxygen diffusivity (D_C_).

### Data analysis

#### Demographic and clinical data

Independent sample t-tests were run between groups for each of the demographic, cognitive and behavioural measures. For demographics such as sex, a chi squared (X^2^) test was run.

#### MRI data

Our analysis aimed to extract cerebral physiological parameters, from dual-calibrated functional MRI (dc-fMRI), representative of whole GM and regional GM in PwMS and controls. Accurate tissue identification required lesion filling in PwMS, followed by tissue segmentation. A biophysical model was applied to the BOLD-ASL signals to estimate physiological parameters voxel-wise. We restricted our subsequent analysis of physiology to voxels containing mainly GM in order to reduce the possibility of any observed differences between PwMS and controls being driven by GM tissue loss in the PwMS. We conducted a region-wise analysis of physiological parameters to further reduce sensitivity to GM partial volume errors associated with tissue loss and to enhance sensitivity through regional averaging. All MRI data were pre-processed and analysed using an in-house written MATLAB (R2015a) pipeline and FSL (Jenkinson, Beckmann, Behrens, Woolrich, & Smith, 2012). Segmentation of GM and WM was performed with FSL-FAST (Zhang, Brady, & Smith, 2001). Registrations were carried out with FSL-FLIRT (S. Smith et al., 2001) and non-brain removal with BET (S. M. Smith, 2002).

#### Lesion filling

T2, PD and FLAIR images were brain extracted. The T2-FLAIR image was transformed to the same space as T2-PD image. Using *JIM* (Version 6.0), lesions where defined manually using the contour ROI tool, without 3D propagation. All three image contrasts were used to locate the lesions. This lesion map was exported as a NIFTI file, with a pixel area threshold of 50%, and a total lesion volume was calculated for each PwMS. For the lesion filling of the T1 weighted image, the PD image was registered to the T1 image, and the lesion map was binarised and transformed to T1-space. The lesion map was thresholded at 0.4 to approximately preserve the size of the original lesion map after transformation and to allow a small amount of inflation, in case of registration errors. This map was then binarised. The function *lesion_filling* (Battaglini, Jenkinson, & De Stefano, 2012) was used: this fills the lesion area with intensities similar to those of the non-lesion neighbourhood.

#### GM and WM volumes

Using the lesion-filled T1 image, FSL-FAST was run to investigate for differences in GM and WM volume between groups. GM, WM and CSF partial volume estimates (PVE) were created through FAST (FSL tool) segmentation of the T1-weighted image. A probability threshold of 0.5 was used to identify GM and WM voxels for each participant. t-tests were carried out using RStudio (http://www.rstudio.com/) in order to investigate between-group differences in global WM and GM volumes. In addition, we calculated the number of GM voxels in each brain region, as defined in the regional analysis (below), using the partial volume image, set with a GM partial volume threshold of above 0.5. Overall, this analysis allowed us to establish whether possible GM volume differences contribute to differences in vascular/metabolic function in the PwMS compared to the controls.

#### dc-fMRI modelling

Analysis of the dc-fMRI data were conducted using a Frequency-Domain Machine Learning (FML) method (Germuska et al., 2020). The method takes advantage of the speed and noise resilience of artificial neural networks to make rapid and robust parameter estimates without the need to apply spatial smoothing. Once the FML pipeline had been run for each participant, the output maps (CBF ml/100 g/m, OEF, CMRO_2_ μmol/100 g/m) were analysed in each subject’s dc-functional MRI space. Functional data (TE1, first echo) was registered to the structural T1-weighted image using FSL’s *epi-reg* tool. Using the inverse of this matrix, the T1-image and the GM-PVE image were transformed to functional space. The GM-PVE image (thresholded at 0.5, as explained before) was binarised and used as a GM mask for calculating whole-GM and regional-GM parameter estimates from the FML output maps (median values). From here, statistical analysis was run to identify group differences in CBF, OEF and CMRO_2_. D_C_ (μmol/100g/mmHg/min) was calculated using GM median values of OEF and CMRO_2_, assuming minimal oxygen tension at the mitochondria. Independent sample t-tests were used to test for differences between patients and controls in CBF, OEF, CMRO_2_, and D_C._ All the reported *p-*values are two-tailed.

#### Regional analysis

Registration matrices were defined to transform dc-fMRI space data to MNI space, using FSL’s FLIRT (linear) function. Registration between native and standard space involved registering the T1 structural image to MNI space using a standard template. The inverse of this matrix for each participant was used to transform the automated anatomical labelling (AAL) atlas to native dc-fMRI space (Tzourio-Mazoyer et al., 2002), where vascular/metabolic parameters were extracted for each region. The AAL atlas includes 116 regions across the cortex and cerebellum. Due to the limited number of slices in our dc-fMRI acquisition, it was not possible to include the cerebellum in the analysis, leaving 83 regions. Median values for each parameter (CBF, OEF, and CMRO_2_) were extracted from each region with GM PV threshold of 0.5. Regional D_C_ values were calculated using the median values for regional OEF and CMRO_2_ (Germuska et al., 2019). Statistical analysis involved running t-tests between patients and controls for each region (FDR corrected) using custom R and Python scripts. All *p-*values reported are two-tailed.

#### Partial volume estimate analysis

To assess whether differences in the physiological parameters vary with the proportion of GM present in the voxels, we ran a partial volume estimate (PVE) analysis. Binarised masks were created from the GM PVE in individual subject functional space for different GM PVE ranges (7 bins of equal width from 0.1-0.8). Within each mask for each parameter (CBF, OEF and CMRO_2_), median values were calculated. D_C_ was calculated using median values of OEF and CMRO_2_ in each GM PVE range. A mixed ANOVA was run to look at main effects (GM PVE bin; Group) and interaction effects (GM PVE bin*Group). All the reported *p-* values are two-tailed.

## Results

### Clinical, demographics, and conventional MRI characteristics

Twenty-two PwMS and 20 controls were included in the study. Out of the 22 patients included in the final analysis, 11 were on disease modifying drugs (DMD) at the time of testing. Mean disease duration was 8.1±1.04 years. Significant group differences were seen for the timed 25-foot walk (*p*=0.006). No other significant group differences were observed (**Table 1**). Brain volume analysis showed no significant differences in GM [*t*(38.87) = 1.05, *p*=0.302] or WM [*t*(34.36) = 1.90, *p*=0.065] volume between groups (**Table 1**). Blood hemoglobin (g/L) content was not significantly different between patients (135.7 ± 3.29 g/L) and controls (136.5 ± 2.83 g/L), *t*(40) = 0.19, *p*=0.849).

**Table 1.** Demographic and clinical characteristics for patients and controls. Values are mean ± SD, unless stated otherwise. All group differences are tested with t-tests, excluding sex, which was tested with chi-squared. *Abbreviations:* EDSS= Expanded Disability Status Scale; MSIS = Multiple Sclerosis Impact Scale; 9HPT-DH = 9 Hole Peg Test – Dominant Hand; T25FW = Timed 25 foot walk; PASAT = Paced Auditory Serial Addition Test.

|  | **Patients** | **Controls** | **p** |
| --- | --- | --- | --- |
| Age | 35.55 ± 7.24 | 34.15 ± 5.69 | 0.489 |
| Sex (M/F) | 5/17 | 6/14 | 0.854 |
| Education (years) | 17.14 ± 1.96 | 18.58 ± 2.81 | 0.699 |
| Disease duration (years) | 8.20 ± 4.85 | - | - |
| Number of patients on DMT's | 11 | - | - |
| Visual Acuity-L(cumulative) | 3.46 ± 1.17 | 3.53 ± 3.06 | 0.928 |
| Visual Acuity-R (cumulative) | 3.25 ± 1.51 | 2.88 ± 1.32 | 0.402 |
| 9HPT-dominant (seconds) | 20.98 ± 4.90 | 20.18 ± 2.28 | 0.235 |
| 9HPT-non-dominant (seconds) | 22.92 ± 2.55 | 21.25 ± 2.97 | 0.058 |
| T25-FW (seconds) | 5.00 ± 1.49 | 3.98 ± 0.63 | 0.006 |
| PASAT-3s (correct/60) | 44.50 ± 12.91 | 48.10 ± 10.54 | 0.326 |
| PASAT-2s (correct/60) | 34.05 ± 11.74 | 35.90 ± 12.21 | 0.623 |
| SDMT | 59.14 ± 7.47 | 62.40 ± 9.76 | 0.499 |
| T2 hyperintense lesion volume (mm<sup>3</sup>) | 10148.38 ± 9,094.02 | - | - |
| White matter volume (mm<sup>3</sup>) | 509,916.2 ± 46148.72 | 542,906 ± 63737.52 | 0.065 |
| Grey matter volume (mm<sup>3</sup>) | 530,891 ± 44,462.88 | 545,840.8 ± 47,835.55 | 0.302 |

### Between group differences in GM physiology

Patients had significantly lower global CBF [PwMS vs. controls: 44.91 ± 6.10 vs. 48.90 ± 5.87 ml/100g/min, *t*(40) = 2.70, *p*=0.010], global CMRO_2_ [PwMS vs. controls: 117.69 ± 17.31 vs. 136.49 ± 14.48 μmol/100g/min, *t*(40) = 3.80, *p* <.001] and D_C_ [PwMS vs. controls: 2.70 ± 0.51 vs. 3.18 ± 0.41 μmol/100g/mmHg/min, *t*(40) = 3.32, *p*=0.002] compared to controls (**Figure 2**). There were no significant group differences in global OEF [PwMS vs. controls: 0.38 ± 0.09 vs. 0.39 ± 0.02, *t*(40) = 0.93, *p* = 0.358] (**Figure 2**). Results of the regional analysis revealed significant widespread reductions in CMRO_2_ and D_C_ in PwMS compared to controls (*p* < 0.05, FDR corrected) (**Figure 3**, **Table 2**). No significant group differences were seen for CBF or OEF across any of the regions. None of the data violated the assumptions associated with independent samples t-tests.

**Figure 2.**
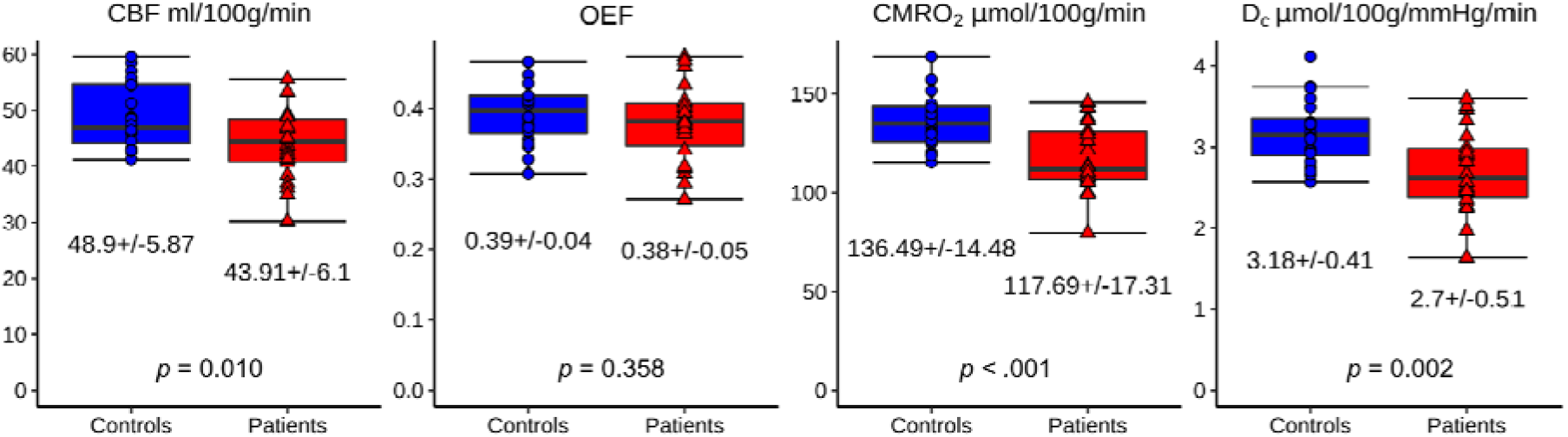
CBF, OEF, CMRO_2_ and D_C_ in patients (red triangles) and in controls (blue circles). Results reflect median values in the GM and then mean across participants (PVE threshold of 0.5). There was a significant difference between groups in CBF, CMRO_2_ and D_C_, but not in OEF. *Abbreviations:* Grey matter (GM) cerebral blood flow (CBF), oxygen extraction fraction (OEF), cerebral metabolic rate of oxygen consumption (CMRO_2_) and capillary oxygen diffusivity (D_C_).

**Figure 3.**
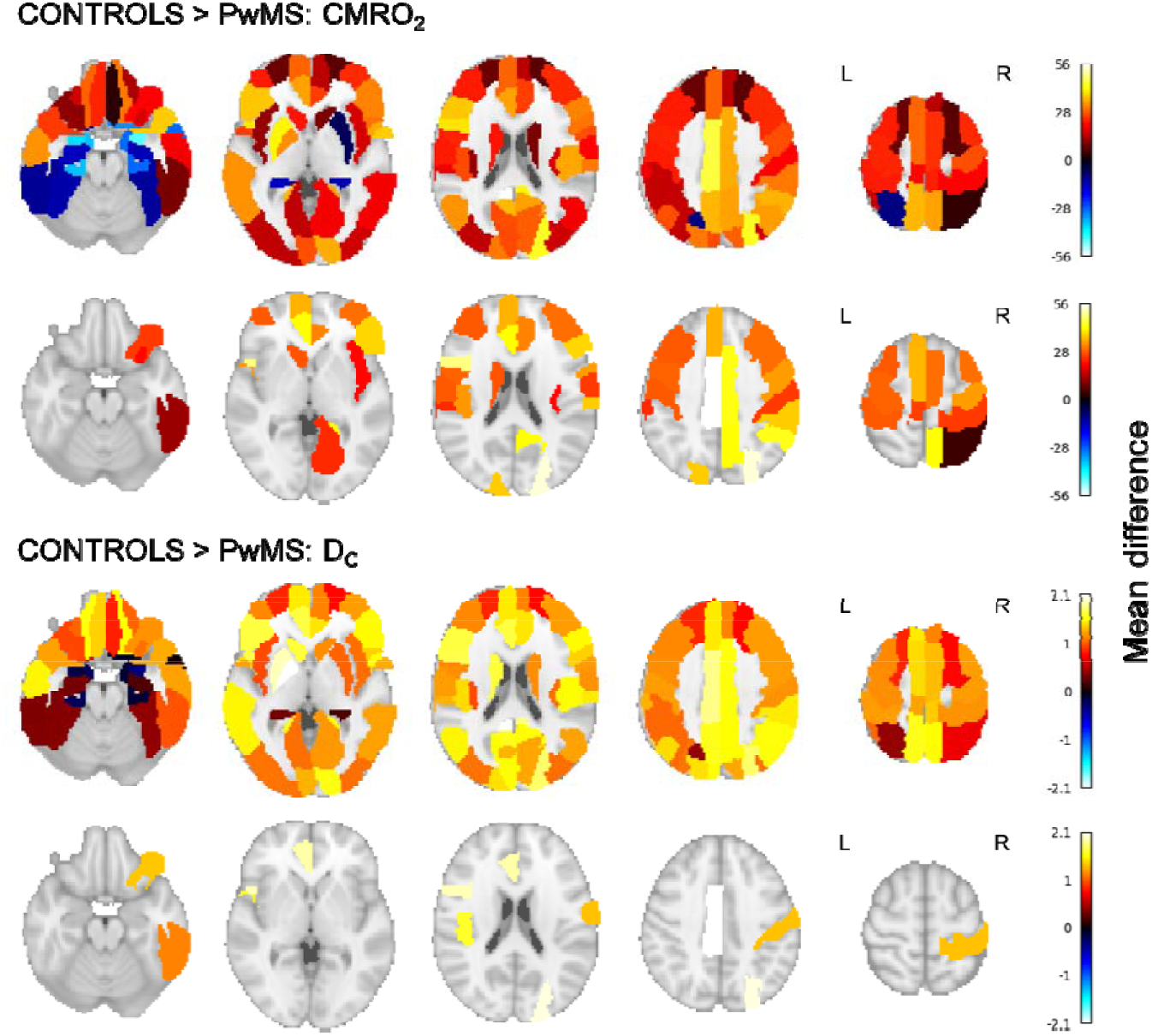
Unthresholded (top row) and thresholded (bottom row) group mean differences (controls>PwMS) for each regional CMRO_2_ (μmol/100g/min) and D_C_ (μmol/100g/mmHg/min) parameters. Differences between groups are reported by the colour-coded scale. Warm colours indicate regions where controls have greater values than patients; cold colours indicate regions where patients have greater values than controls. Reductions in CMRO_2_ and D_C_ appear to be widespread in patients. *Abbreviations:* Cerebral metabolic rate of oxygen consumption (CMRO_2_) and capillary oxygen diffusivity (D_C_).

**Table 2.** Results from the regional analysis reporting the areas in which significant reductions were observed for patients compared to controls. There were only significant regional differences found for CMRO_2_ and D_C_. Analysis includes FDR corrected results only. *Abbreviations:* cerebral metabolic rate of oxygen consumption (CMRO_2_) and capillary oxygen diffusivity (D_C_).

| Region | Parameter | t-statistic | p-value (FDR corr) |
| --- | --- | --- | --- |
| Cingulum_Mid_L | CMRO<sub>2</sub> | 4.06 | 0.019 |
| Frontal_Inf_Orb_R | CMRO<sub>2</sub> | 3.55 | 0.027 |
| Occipital_Sup_R | CMRO<sub>2</sub> | 3.56 | 0.027 |
| Postcentral_R | CMRO<sub>2</sub> | 3.40 | 0.027 |
| Temporal_Inf_R | CMRO<sub>2</sub> | 3.38 | 0.027 |
| Frontal_Inf_Oper_L | CMRO<sub>2</sub> | 3.31 | 0.027 |
| Precentral_R | CMRO<sub>2</sub> | 3.17 | 0.030 |
| Rolandic_Oper_L | CMRO<sub>2</sub> | 3.08 | 0.030 |
| Cingulum_Ant_L | CMRO<sub>2</sub> | 3.24 | 0.030 |
| Cingulum_Ant_R | CMRO<sub>2</sub> | 3.04 | 0.030 |
| Postcentral_L | CMRO<sub>2</sub> | 3.08 | 0.030 |
| Caudate_L | CMRO<sub>2</sub> | 3.04 | 0.030 |
| Precuneus_R | CMRO<sub>2</sub> | 2.97 | 0.031 |
| SupraMarginal_R | CMRO<sub>2</sub> | 2.87 | 0.039 |
| Parietal_Sup_R | CMRO<sub>2</sub> | 2.82 | 0.043 |
| Supp_Motor_Area_L | CMRO<sub>2</sub> | 2.74 | 0.045 |
| Occipital_Sup_L | CMRO<sub>2</sub> | 2.76 | 0.045 |
| Precentral_L | CMRO<sub>2</sub> | 2.69 | 0.045 |
| Parietal_Inf_R | CMRO<sub>2</sub> | 2.70 | 0.045 |
| Frontal_Mid_L | CMRO<sub>2</sub> | 2.65 | 0.046 |
| Lingual_R | CMRO<sub>2</sub> | 2.65 | 0.046 |
| Frontal_Mid_R | CMRO<sub>2</sub> | 2.62 | 0.046 |
| Frontal_Inf_Tri_R | CMRO<sub>2</sub> | 2.57 | 0.047 |
| Supp_Motor_Area_R | CMRO<sub>2</sub> | 2.52 | 0.047 |
| Frontal_Sup_Medial_L | CMRO<sub>2</sub> | 2.53 | 0.047 |
| Insula_R | CMRO<sub>2</sub> | 2.56 | 0.047 |
| SupraMarginal_L | CMRO<sub>2</sub> | 2.53 | 0.047 |
| Paracentral_Lobule_L | CMRO<sub>2</sub> | 2.57 | 0.047 |
| Cingulum_Mid_R | CMRO<sub>2</sub> | 2.49 | 0.049 |
| Cingulum_Mid_L | D<sub>c</sub> | 3.83 | 0.036 |
| Cingulum_Ant_L | D<sub>c</sub> | 3.23 | 0.045 |
| Frontal_Inf_Oper_L | D<sub>c</sub> | 3.04 | 0.045 |
| Frontal_Inf_Orb_R | D<sub>c</sub> | 3.08 | 0.045 |
| Occipital_Sup_R | D<sub>c</sub> | 3.45 | 0.045 |
| Postcentral_R | D<sub>c</sub> | 3.02 | 0.045 |
| Rolandic_Oper_L | D<sub>c</sub> | 3.21 | 0.045 |
| Temporal_Inf_R | D<sub>c</sub> | 3.26 | 0.045 |

### Relationship between brain physiology and measures of damage and disability

There was a significant correlation between white matter T2 hyperintense lesion volume and global GM CBF (*R* = −0.52, *p* = 0.014), as well as between white matter T2 hyperintense lesion volume and global GM CBF and global GM CMRO_2_ (*R* = −0.48, *p* = 0.023). There were no significant correlations between T2 lesional damage and global OEF and D_c_. Critically, no significant correlations were observed between EDSS scores and CBF (*R* = −.36, *p* = 0.103), OEF (*R* = −.12, *p* = 0.606), CMRO_2_ (*R* = −0.06, *p* = 0.781) or D_C_ (*R*=0.16, *p* = 0.492). Similarly no significant correlations between regional vascular/metabolic parameters (CBF; OEF; CMRO2; DC) and other measures of disability (MSIS; SDMT; PASAT; visual acuity or 25-foot walk) were observed after correction for multiple comparisons (*p* > 0.05, FDR corrected).

### The effect of GM partial volume on physiological parameter estimates

The partial volume analysis of global GM revealed a significant main effect of PVE bin for all parameters (**Figure 4** and **Table 3**): CBF (β = 4.76, *p* < 0.001), OEF (β = 0.01, *p* < 0.001), CMRO_2_ (β *=* 13.92, *p* = 0.001), and D_C_ (β = 0.35, *p* <0.001), broadly indicating an increase in the estimate of each parameter with increasing proportion of GM. We also observed a significant main effect of group for CBF (β = −3.24, *p* = 0.019), CMRO_2_ (β = −11.18, *p* = 0.004), and D_C_ (β = −0.27, *p* = 0.013), but not OEF (β = −0.02, *p* = 0.118), suggesting consistently lower values of CBF, CMRO_2_ and D_C_ across different GM thresholds in PwMS compared to controls. There were no significant interactions PVE bin*Group for any of the parameters [CBF (β = −0.34, *p* = 0.263), OEF (β = 0.00, *p* = 0.788), CMRO_2_ (β = −1.47, *p* = 0.089), D_C_ (β = −0.04, *p* = 0.101)], indicating that GM thresholds have little influence on detecting physiological differences between PwMS and controls. To evaluate whether regional GM volume may have contributed to the regional group differences of physiological parameters, we calculated the number of voxels (PVE threshold >0.5) in each ROI and compared this between groups. Results showed no significant differences between PwMS and controls in any of the AAL regions (*p* > 0.05, FDR corrected).

**Figure 4.**
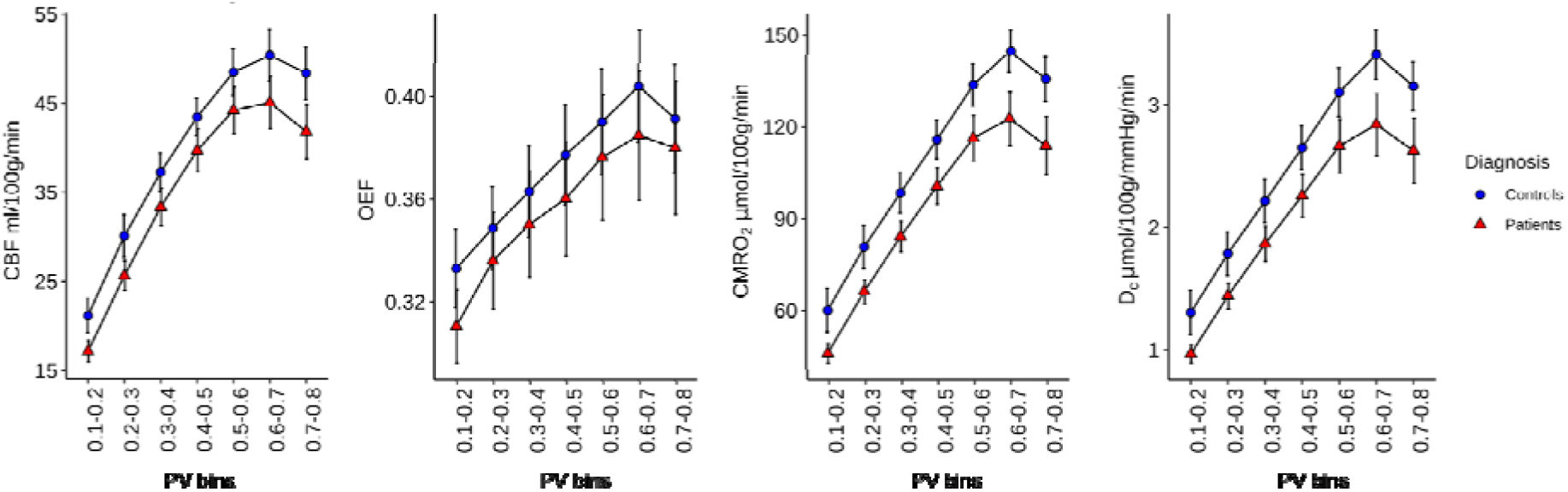
Graphs showing the average GM for each physiological parameter (CBF, OEF, CMRO_2_, and D_C_) across different PVE bins for PwMS (red) and controls (blue). Bars represent 95% confidence intervals. *Abbreviations:* cerebral blood flow (CBF), oxygen extraction fraction (OEF), cerebral metabolic rate of oxygen consumption (CMRO_2_) and capillary oxygen diffusivity (D_C_).

**Table 3.**
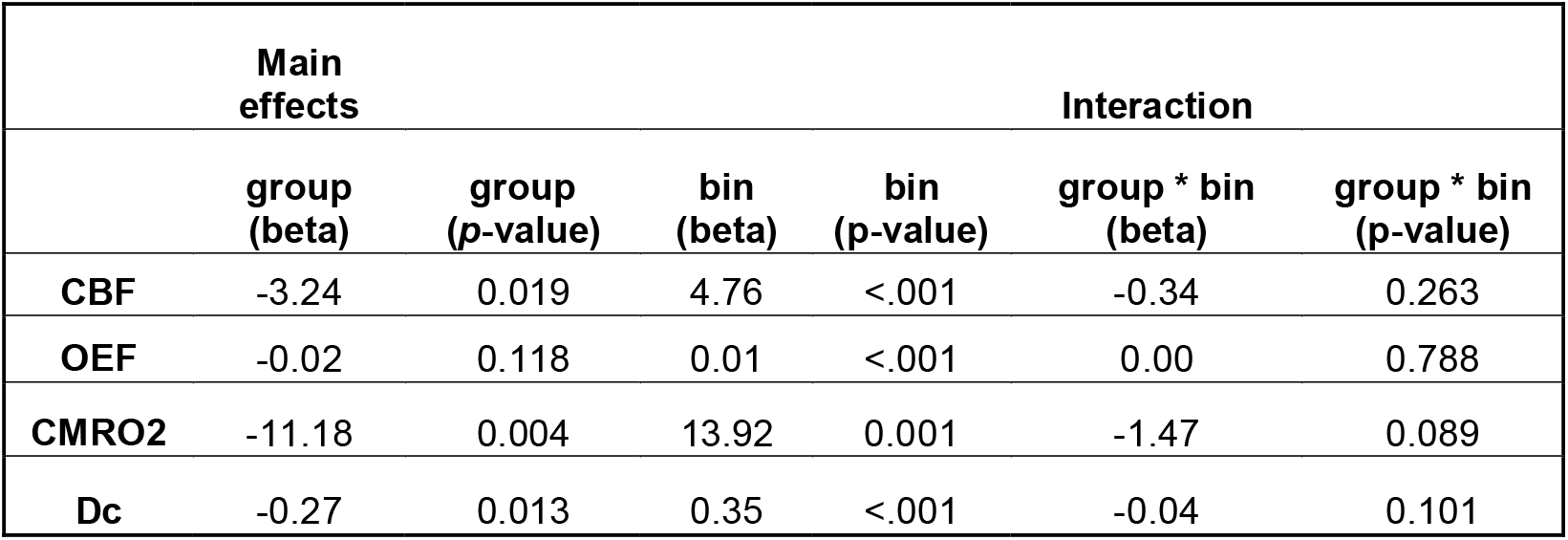
Results of partial volume analysis. Table shows main effect and interaction effects between Group (PwMS vs. controls) and partial volume bin. *Abbreviations:* cerebral blood flow (CBF), oxygen extraction fraction (OEF), cerebral metabolic rate of oxygen consumption (CMRO_2_) and capillary oxygen diffusivity (D_C_).

## Discussion

This study quantifies brain O_2_ supply and consumption in PwMS using a newly developed multiparametric method, the dc-fMRI (Germuska et al., 2019; Germuska et al., 2016). We observed reductions in CBF, CMRO_2_, and D_C_ within the GM of PwMS compared to matched healthy controls, but no between-group difference was observed for OEF. Regional analysis confirmed statistically significant reductions in GM CMRO_2_ and D_C_ only. Overall, these findings suggest vascular and metabolic dysregulation in MS. The lack of association between the quantity of GM and the physiological parameters at the regional level, along with the absence of any significant group differences in GM volume, suggests that the described changes in tissue physiology may precede significant GM tissue loss in PwMS.

### Changes in brain physiology in MS

The observed reductions in GM CBF and CMRO_2_ may reflect (i) microvascular inefficiency in transferring O_2_ from the capillary to the mitochondria, (ii) reduced demand for nutrients including O_2_, (iii) an inability of the tissue to use metabolic substrates for energy release or (iv) primary loss of blood supply at the macrovascular level. The latter explanation is unlikely, given that Dc is reduced, but OEF is not elevated, suggesting that the reduction in Dc is associated with local impairment of O_2_ diffusion at the capillary bed rather than with global change in blood supply (Fan et al., 2015a; Ge et al., 2012; Watchmaker et al., 2018). The global and localized reduction of effective O_2_ diffusivity of the capillary network (D_C_) suggests an impairment of microvascular function, the specific nature of which requires further elucidation. Reduced microvascular density has been observed in MS (Lanzillo et al., 2019). Emerging imaging approaches including optical coherence tomography angiography (OCTA), which permits mapping of microvasculature across the retina, has revealed that PwMS have a reduced density of vessels in superficial and deep vascular plexuses, correlating with function and MS disability scores (Murphy et al., 2020; Wang, Murphy, Caldito, Calabresi, & Saidha, 2018). We suggest that a reduced capillary density may lead to an increase in vascular resistance, which would result in a reduction in flow and a limited O_2_ exchange. Destabilization of the blood brain barrier (BBB) and damage to pericytes as a result of neuroinflammatory processes may create a potential pathological pathway for microvascular dysregulation (Brown et al., 2019; Cheng et al., 2018; Marshall et al., 2014).

The reduced rate of CMRO_2_ may reflect histotoxic hypoxia that is an increased energy demand of impulse conduction along excitable demyelinated axons accompanied by a failure in energy production with intact O_2_ supply. Histotoxic hypoxia has been identified in MS and referred to as virtual hypoxia (Trapp & Stys, 2009). Altered energy production underlying virtual hypoxia may be due to soluble mediators of inflammation that impair mitochondrial function (Mahad et al 2009) and/or to oxidative damage to mitochondrial DNA that may further disrupt mitochondrial activity (Lu et al 2000), ultimately leading to cortical degeneration (Haider et al 2016). Diffuse hypoxia is likely to be present in MS and may be a critical pathological mechanism in MS, as the hypoxic and inflammatory responses are intimately linked (Yang & Dunn 2019): hypoxia exacerbates the inflammatory response (Snyder, Shell, Cunningham, & Cunningham, 2017) and inflammation can damage the vascular endothelium, reduce vasoreactivity and promote the influx of leucocytes. In acute inflammation, an increased metabolic demand from inflammatory cells may also increase oxidative stress and exacerbate hypoxia. This cascade of events may chronically lead to further tissue damage and disability.

Regional analysis confirmed CMRO_2_ and D_C_ reductions to be widespread across the cortex, but with the most significant reductions located around the cingulate and frontal cortex (Eshaghi et al., 2018). Typically, MS lesions are localised particularly in periventricular WM which would significantly affect the tracts connecting key frontal and cingulate regions, and thus may make these regions more susceptible to degeneration (Charil et al., 2007; Haider et al., 2016).

We considered the possibility that partial volume differences between the groups might bias our results and compared physiological estimates as a function of GM partial volume. We only included bins up to a GM partial volume threshold of 0.7-0.8 as there is a limited number of voxels from which to sample for higher thresholds applied to some anatomical regions. As expected, estimated values of physiological parameters increased as the proportion of GM increased. The CBF, CMRO_2_, and D_C_ differences between PwMS and controls were robust to GM PV variation, suggesting that the observed group differences reflects aspects of tissue physiology without being biased by tissue loss.

### Relationship between brain physiology, damage and disability

Our results demonstrated that increasing levels of T2 lesion volume is associated with reduced CBF and CMRO_2_ in GM, consistently with previous findings (Ge et al., 2012). It could be assumed that a higher WM lesion load reflects a higher GM lesion load (Bodini et al., 2009; Muhlau et al., 2013; Steenwijk et al., 2016). However, the relationship is not clearly established (Calabrese et al., 2013). Future studies looking at GM physiology in MS could benefit from the inclusion of scans more sensitive to GM damage and myelination (Abidi, Faeghi, Mardanshahi, & Mortazavi, 2017).

Our data indicated no association between disability scales and tissue physiological status. Lack of direct correlations between behavioural and physiological measures may arise from the presence of adaptation to underlying tissue dysfunction and damage that modulates behavioural performance. While adaptation may confound the relationship between disability measures and brain physiology, we speculate that tissue physiology may be predictive of future tissue damage and behavioural alterations, an avenue that remains to be explored.

### Potential limitations of the study

Our results should be considered in the light of the following limitations. Given that our calibrated fMRI method relies on arterial spin labelling based measurements of CBF we are only able to reliably estimate physiological parameters in GM. This limits the comparisons we can make with other studies that summarise O_2_ metabolic changes across GM and WM (Fan et al., 2015b; Ge et al., 2012; West et al., 2020). Regional differences in the significance of CMRO_2_ and D_C_ reductions may arise not only from disease effects, but also from variation in the sensitivity of the technique that is dependent on regional within- and between-subject variation of BOLD and CBF measurements. The scans we used to identify lesions are not generally sensitive to lesions within GM. Therefore, our reported associations with GM physiology are limited to WM lesional damage.

## Conclusions

Our findings suggest that a metabolic dysregulation occurs in patients MS, resulting in reduced O_2_ supply and utilisation. These changes may precede GM tissue loss, thereby providing potentially promising avenues for neuroprotective interventions. The use of multi-parametric dc-MRI approach can provide valuable quantitative data regarding altered brain physiology in MS and, with a regional approach, may help identify target areas for early identification of tissue dysfunction and its subsequent evolution.

## Acknowledgments

This project was supported by an EPSRC grant and a Wellcome PhD studentship. HLC and MG are funded by a Wellcome Strategic Award [104943/Z/14/Z]. We would like to thank the patients with MS and their families, along with the healthy volunteers for their time and support that made this research possible.

## Contributions

Study design (HLC, RS, CF, RGW, VT); data collection (HLC, RS, EP, NS, SK); data analysis (HLC, MG, TML, RS, SBD); writing manuscript (HLC, RGW, VT).

## References

Abidi, Z., Faeghi, F., Mardanshahi, Z., & Mortazavi, H. (2017). Assessment of the diagnostic accuracy of double inversion recovery sequence compared with FLAIR and T2W_TSE in detection of cerebral multiple sclerosis lesions. Electron Physician, 9(4), 4162–4170. doi:10.19082/4162

Battaglini, M., Jenkinson, M., & De Stefano, N. (2012). Evaluating and reducing the impact of white matter lesions on brain volume measurements. Human Brain Mapping, 33(9), 2062–2071. doi:10.1002/hbm.21344

Benedict, R. H. B., DeLuca, J., Phillips, G., LaRocca, N., Hudson, L. D., Rudick, R., & Assessm, M. S. O. (2017). Validity of the Symbol Digit Modalities Test as a cognition performance outcome measure for multiple sclerosis. Multiple Sclerosis Journal, 23(5), 721–733. doi:10.1177/1352458517690821

Bodini, B., Khaleeli, Z., Cercignani, M., Miller, D. H., Thompson, A. J., & Ciccarelli, O. (2009). Exploring the relationship between white matter and gray matter damage in early primary progressive multiple sclerosis: an in vivo study with TBSS and VBM. Human Brain Mapping, 30(9), 2852–2861. doi:10.1002/hbm.20713

Brooks, D. J., Leenders, K. L., Head, G., Marshall, J., Legg, N. J., & Jones, T. (1984). Studies on regional cerebral oxygen utilisation and cognitive function in multiple sclerosis. J Neurol Neurosurg Psychiatry, 47(11), 1182–1191. doi:10.1136/jnnp.47.11.1182

Brown, L. S., Foster, C. G., Courtney, J. M., King, N. E., Howells, D. W., & Sutherland, B. A. (2019). Pericytes and Neurovascular Function in the Healthy and Diseased Brain. Frontiers in Cellular Neuroscience, 13, 282. doi:10.3389/fncel.2019.00282

Bulte, D. P., Kelly, M., Germuska, M., Xie, J., Chappell, M. A., Okell, T. W., … Jezzard, P. (2012). Quantitative measurement of cerebral physiology using respiratory-calibrated MRI. Neuroimage, 60(1), 582–591. doi:10.1016/j.neuroimage.2011.12.017

Calabrese, M., Romualdi, C., Poretto, V., Favaretto, A., Morra, A., Rinaldi, F., … Gallo, P. (2013). The changing clinical course of multiple sclerosis: a matter of gray matter. Annals of Neurology, 74(1), 76–83. doi:10.1002/ana.23882

Charil, A., Dagher, A., Lerch, J. P., Zijdenbos, A. P., Worsley, K. J., & Evans, A. C. (2007). Focal cortical atrophy in multiple sclerosis: Relation to lesion load and disability. Neuroimage, 34(2), 509–517. doi:10.1016/j.neuroimage.2006.10.006

Cheng, J., Korte, N., Nortley, R., Sethi, H., Tang, Y., & Attwell, D. (2018). Targeting pericytes for therapeutic approaches to neurological disorders. Acta Neuropathologica, 136(4), 507–523. doi:10.1007/s00401-018-1893-0

Cutter, G. R., Baier, M. L., Rudick, R. A., Cookfair, D. L., Fischer, J. S., Petkau, J., … Willoughby, E. (1999). Development of a multiple sclerosis functional composite as a clinical trial outcome measure. Brain, 122 (Pt 5), 871–882. doi:10.1093/brain/122.5.871

Davies, A. L., Desai, R. A., Bloomfield, P. S., McIntosh, P. R., Chapple, K. J., Linington, C., … Smith, K. J. (2013). Neurological deficits caused by tissue hypoxia in neuroinflammatory disease. Annals of Neurology, 74(6), 815–825. doi:10.1002/ana.24006

Debernard, L., Melzer, T. R., Van Stockum, S., Graham, C., Wheeler-Kingshott, C. A. M., Dalrymple-Alford, J. C., … Mason, D. F. (2014). Reduced grey matter perfusion without volume loss in early relapsing-remitting multiple sclerosis. Journal of Neurology Neurosurgery and Psychiatry, 85(5), 544–551. doi:10.1136/jnnp-2013-305612

Eshaghi, A., Marinescu, R. V., Young, A. L., Firth, N. C., Prados, F., Jorge Cardoso, M., … Ciccarelli, O. (2018). Progression of regional grey matter atrophy in multiple sclerosis. Brain, 141(6), 1665–1677. doi:10.1093/brain/awy088

Fan, A. P., Govindarajan, S. T., Kinkel, R. P., Madigan, N. K., Nielsen, A. S., Benner, T., … Mainero, C. (2015a). Quantitative oxygen extraction fraction from 7-Tesla MRI phase: reproducibility and application in multiple sclerosis. Journal of Cerebral Blood Flow and Metabolism, 35(1), 131–139. doi:10.1038/jcbfm.2014.187

Fan, A. P., Govindarajan, S. T., Kinkel, R. P., Madigan, N. K., Nielsen, A. S., Benner, T., … Mainero, C. (2015b). Quantitative oxygen extraction fraction from 7-Tesla MRI phase: reproducibility and application in multiple sclerosis. J Cereb Blood Flow Metab, 35(1), 131–139. doi:10.1038/jcbfm.2014.187

Gauthier, C. J., & Hoge, R. D. (2013). A generalized procedure for calibrated MRI incorporating hyperoxia and hypercapnia. Human Brain Mapping, 34(5), 1053–1069. doi:10.1002/hbm.21495

Ge, Y., Zhang, Z., Lu, H., Tang, L., Jaggi, H., Herbert, J., … Grossman, R. I. (2012). Characterizing brain oxygen metabolism in patients with multiple sclerosis with T2-relaxation-under-spin-tagging MRI. J Cereb Blood Flow Metab, 32(3), 403–412. doi:10.1038/jcbfm.2011.191

Germuska, M., Chandler, H., Okell, T., Fasano, F., Tomassini, V., Murphy, K., & Wise, R. (2020). A frequency-domain machine learning method for dual-calibrated fMRI mapping of oxygen extraction fraction (OEF) and cerebral metabolic rate of oxygen consumption (CMRO2). Front Artif Intell, 3. doi:10.3389/frai.2020.00012

Germuska, M., Chandler, H. L., Stickland, R. C., Foster, C., Fasano, F., Okell, T. W., … Wise, R. G. (2019). Dual-calibrated fMRI measurement of absolute cerebral metabolic rate of oxygen consumption and effective oxygen diffusivity. Neuroimage, 184, 717–728. doi:10.1016/j.neuroimage.2018.09.035

Germuska, M., Merola, A., Murphy, K., Babic, A., Richmond, L., Khot, S., … Wise, R. G. (2016). A forward modelling approach for the estimation of oxygen extraction fraction by calibrated fMRI. Neuroimage, 139, 313–323. doi:10.1016/j.neuroimage.2016.06.004

Germuska, M., & Wise, R. G. (2019). Calibrated fMRI for mapping absolute CMRO2: Practicalities and prospects. Neuroimage, 187, 145–153. doi:10.1016/j.neuroimage.2018.03.068

Haider, L., Zrzavy, T., Hametner, S., Hoftberger, R., Bagnato, F., Grabner, G., … Lassmann, H. (2016). The topograpy of demyelination and neurodegeneration in the multiple sclerosis brain. Brain, 139(Pt 3), 807–815. doi:10.1093/brain/awv398

Hobart, J., Lamping, D., Fitzpatrick, R., Riazi, A., & Thompson, A. (2001). The Multiple Sclerosis Impact Scale (MSIS-29): a new patient-based outcome measure. Brain, 124(Pt 5), 962–973. doi:10.1093/brain/124.5.962

Hoge, R. D. (2012). Calibrated FMRI. Neuroimage, 62(2), 930–937. doi:10.1016/j.neuroimage.2012.02.022

Jenkinson, M., Beckmann, C. F., Behrens, T. E., Woolrich, M. W., & Smith, S. M. (2012). Fsl. Neuroimage, 62(2), 782–790. doi:10.1016/j.neuroimage.2011.09.015

Kurtzke, J. F. (1983). Rating neurologic impairment in multiple sclerosis: an expanded disability status scale (EDSS). Neurology, 33(11), 1444–1452. doi:10.1212/wnl.33.11.1444

Lanzillo, R., Cennamo, G., Moccia, M., Criscuolo, C., Carotenuto, A., Frattaruolo, N., … Brescia Morra, V. (2019). Retinal vascular density in multiple sclerosis: a 1-year follow-up. European Journal of Neurology, 26(1), 198–201. doi:10.1111/ene.13770

Law, M., Saindane, A. M., Ge, Y., Babb, J. S., Johnson, G., Mannon, L. J., … Grossman, R. I. (2004). Microvascular abnormality in relapsing-remitting multiple sclerosis: perfusion MR imaging findings in normal-appearing white matter. Radiology, 231(3), 645–652. doi:10.1148/radiol.2313030996

Lublin, F. D., Reingold, S. C., Cohen, J. A., Cutter, G. R., Sorensen, P. S., Thompson, A. J., … Polman, C. H. (2014). Defining the clinical course of multiple sclerosis: the 2013 revisions. Neurology, 83(3), 278–286. doi:10.1212/WNL.0000000000000560

Marshall, O., Chawla, S., Lu, H., Pape, L., & Ge, Y. (2016). Cerebral blood flow modulation insufficiency in brain networks in multiple sclerosis: A hypercapnia MRI study. J Cereb Blood Flow Metab, 36(12), 2087–2095. doi:10.1177/0271678X16654922

Marshall, O., Lu, H., Brisset, J. C., Xu, F., Liu, P., Herbert, J., … Ge, Y. (2014). Impaired cerebrovascular reactivity in multiple sclerosis. Jama Neurology, 71(10), 1275–1281. doi:10.1001/jamaneurol.2014.1668

Merola, A., Germuska, M. A., Murphy, K., & Wise, R. G. (2018). Assessing the repeatability of absolute CMRO2, OEF and haemodynamic measurements from calibrated fMRI. Neuroimage, 173, 113–126. doi:10.1016/j.neuroimage.2018.02.020

Merola, A., Germuska, M. A., Warnert, E. A., Richmond, L., Helme, D., Khot, S., … Wise, R. G. (2017). Mapping the pharmacological modulation of brain oxygen metabolism: The effects of caffeine on absolute CMRO2 measured using dual calibrated fMRI. Neuroimage, 155, 331–343. doi:10.1016/j.neuroimage.2017.03.028

Muhlau, M., Buck, D., Forschler, A., Boucard, C. C., Arsic, M., Schmidt, P., … Ilg, R. (2013). White-matter lesions drive deep gray-matter atrophy in early multiple sclerosis: support from structural MRI. Mult Scler, 19(11), 1485–1492. doi:10.1177/1352458513478673

Murphy, O. C., Kwakyi, O., Iftikhar, M., Zafar, S., Lambe, J., Pellegrini, N., … Saidha, S. (2020). Alterations in the retinal vasculature occur in multiple sclerosis and exhibit novel correlations with disability and visual function measures. Mult Scler, 26(7), 815–828. doi:10.1177/1352458519845116

Okell, T. W., Chappell, M. A., Kelly, M. E., & Jezzard, P. (2013). Cerebral blood flow quantification using vessel-encoded arterial spin labeling. J Cereb Blood Flow Metab, 33(11), 1716–1724. doi:10.1038/jcbfm.2013.129

Paling, D., Golay, X., Wheeler-Kingshott, C., Kapoor, R., & Miller, D. (2011). Energy failure in multiple sclerosis and its investigation using MR techniques. Journal of Neurology, 258(12), 2113–2127. doi:10.1007/s00415-011-6117-7

Paulesu, E., Perani, D., Fazio, F., Comi, G., Pozzilli, C., Martinelli, V., … Fieschi, C. (1996). Functional basis of memory impairment in multiple sclerosis: a[18F]FDG PET study. Neuroimage, 4(2), 87–96. doi:10.1006/nimg.1996.0032

Penner, I. K., Raselli, C., Stocklin, M., Opwis, K., Kappos, L., & Calabrese, P. (2009). The Fatigue Scale for Motor and Cognitive Functions (FSMC): validation of a new instrument to assess multiple sclerosis-related fatigue. Mult Scler, 15(12), 1509–1517. doi:10.1177/1352458509348519

Schmithorst, V. J., Hernandez-Garcia, L., Vannest, J., Rajagopal, A., Lee, G., & Holland, S. K. (2014). Optimized simultaneous ASL and BOLD functional imaging of the whole brain. Journal of Magnetic Resonance Imaging, 39(5), 1104–1117. doi:10.1002/jmri.24273

Smith, S., Bannister, P. R., Beckmann, C., Brady, M., Clare, S., Flitney, D., … Zhang, Y. Y. (2001). FSL: New tools for functional and structural brain image analysis. Neuroimage, 13(6), S249–S249.

Smith, S. M. (2002). Fast robust automated brain extraction. Human Brain Mapping, 17(3), 143–155. doi:10.1002/hbm.10062

Snyder, B., Shell, B., Cunningham, J. T., & Cunningham, R. L. (2017). Chronic intermittent hypoxia induces oxidative stress and inflammation in brain regions associated with early-stage neurodegeneration. Physiol Rep, 5(9). doi:10.14814/phy2.13258

Steenwijk, M. D., Geurts, J. J., Daams, M., Tijms, B. M., Wink, A. M., Balk, L. J., … Pouwels, P. J. (2016). Cortical atrophy patterns in multiple sclerosis are non-random and clinically relevant. Brain, 139(Pt 1), 115–126. doi:10.1093/brain/awv337

Sun, X., Tanaka, M., Kondo, S., Hirai, S., & Ishihara, T. (1994). Reduced cerebellar blood flow and oxygen metabolism in spinocerebellar degeneration: a combined PET and MRI study. Journal of Neurology, 241(5), 295–300. doi:10.1007/BF00868436

Sun, X., Tanaka, M., Kondo, S., Okamoto, K., & Hirai, S. (1998). Clinical significance of reduced cerebral metabolism in multiple sclerosis: a combined PET and MRI study. Ann Nucl Med, 12(2), 89–94. doi:10.1007/BF03164835

Thompson, A. J., Banwell, B. L., Barkhof, F., Carroll, W. M., Coetzee, T., Comi, G., … Cohen, J. A. (2018). Diagnosis of multiple sclerosis: 2017 revisions of the McDonald criteria. Lancet Neurology, 17(2), 162–173. doi:10.1016/S1474-4422(17)30470-2

Trapp, B. D., & Stys, P. K. (2009). Virtual hypoxia and chronic necrosis of demyelinated axons in multiple sclerosis. Lancet Neurology, 8(3), 280–291. doi:10.1016/S1474-4422(09)70043-2

Tzourio-Mazoyer, N., Landeau, B., Papathanassiou, D., Crivello, F., Etard, O., Delcroix, N., … Joliot, M. (2002). Automated anatomical labeling of activations in SPM using a macroscopic anatomical parcellation of the MNI MRI single-subject brain. Neuroimage, 15(1), 273–289. doi:10.1006/nimg.2001.0978

Wang, L., Murphy, O., Caldito, N. G., Calabresi, P. A., & Saidha, S. (2018). Emerging Applications of Optical Coherence Tomography Angiography (OCTA) in neurological research. Eye Vis (Lond), 5, 11. doi:10.1186/s40662-018-0104-3

Watchmaker, J. M., Juttukonda, M. R., Davis, L. T., Scott, A. O., Faraco, C. C., Gindville, M. C., … Donahue, M. J. (2018). Hemodynamic mechanisms underlying elevated oxygen extraction fraction (OEF) in moyamoya and sickle cell anemia patients. J Cereb Blood Flow Metab, 38(9), 1618–1630. doi:10.1177/0271678X16682509

West, K., Sivakolundu, D., Maruthy, G., Zuppichini, M., Liu, P., Thomas, B., … Rypma, B. (2020). Baseline cerebral metabolism predicts fatigue and cognition in Multiple Sclerosis patients. Neuroimage Clin, 27, 102281. doi:10.1016/j.nicl.2020.102281

Wise, R. G., Harris, A. D., Stone, A. J., & Murphy, K. (2013). Measurement of OEF and absolute CMRO2: MRI-based methods using interleaved and combined hypercapnia and hyperoxia. Neuroimage, 83, 135–147. doi:10.1016/j.neuroimage.2013.06.008

Wuerfel, J., Bellmann-Strobl, J., Brunecker, P., Aktas, O., McFarland, H., Villringer, A., & Zipp, F. (2004). Changes in cerebral perfusion precede plaque formation in multiple sclerosis: a longitudinal perfusion MRI study. Brain, 127(Pt 1), 111–119. doi:10.1093/brain/awh007

Yang, R., & Dunn, J. F. (2015). Reduced cortical microvascular oxygenation in multiple sclerosis: a blinded, case-controlled study using a novel quantitative near-infrared spectroscopy method. Sci Rep, 5, 16477. doi:10.1038/srep16477

Zhang, Y. Y., Brady, M., & Smith, S. (2001). Segmentation of brain MR images through a hidden Markov random field model and the expectation-maximization algorithm. Ieee Transactions on Medical Imaging, 20(1), 45–57. doi:Doi 10.1109/42.906424

